# Lack of intrinsic postzygotic isolation in haplodiploid male hybrids despite high genetic distance

**DOI:** 10.1101/2020.01.08.898957

**Authors:** Emily E. Bendall, Kayla M. Mattingly, Amanda J. Moehring, Catherine R. Linnen

## Abstract

Evolutionary biologists have long been interested in understanding the mechanisms underlying Haldane’s rule. The explanatory theories of dominance and faster-X, which are based on recessive alleles being expressed in the heterogametic sex, have been proposed as common mechanisms. These mechanisms predict that greater hemizygosity leads to both faster evolution and greater expression of intrinsic postzygotic isolation. Under these mechanisms, haplodiploids should evolve and express intrinsic postzygotic isolation faster than diploids because the entire genome is analogous to a sex chromosome. Here, we measure sterility and inviability in hybrids between *Neodiprion pinetum* and *N. lecontei*, a pair of haplodiplopids that differ morphologically, behaviorally, and genetically. We compare the observed isolation to that expected from published estimates of isolation in diploids at comparable levels of genetic divergence. We find that both male and female hybrids are viable and fertile, which is less isolation than expected. We then discuss several potential explanations for this surprising lack of isolation, including alternative mechanisms for Haldane’s rule and a frequently overlooked quirk of haplodiploid genetics that may slow the emergence of complete intrinsic postzygotic isolation in hybrid males. Finally, we describe how haplodiploids, an underutilized resource, can be used to differentiate between mechanisms of Haldane’s rule.

## Introduction

Barriers to gene flow enable species to diverge along independent evolutionary trajectories. For this reason, the evolution of reproductive isolation is a central focus of speciation research. Although there are many different types of reproductive barriers (Dobzhansky 1951; Coyne and Orr 2004), the most impermeable and permanent of these is intrinsic postzygotic isolation (IPI), which is the inability to produce viable, fertile hybrids. At a genetic level, hybrid inviability and sterility are often caused by the accumulation of Bateson-Dobzhansky-Muller incompatibilities (BDMIs) in diverging populations (Bateson 1909; Dobzhansky 1937; Muller 1942). While neutral or beneficial in the parental genomes, negative epistasis among BDMIs in hybrid genomes results in IPI.

In early stages of speciation, sterility or inviability is often restricted to one sex of the hybrid offspring (Coyne and Orr 1989, 1997). When this occurs, it is almost always the heterogametic sex (XY, ZW) that is sterile or inviable, a pattern known as Haldane’s rule (Haldane 1922; Schilthuizen et al. 2011). To date, multiple non-mutually exclusive mechanisms have been proposed to explain Haldane’s rule. Two explanations that have gained considerable empirical support are dominance theory and faster-X theory (Schilthuizen et al. 2011; Delph and Demuth 2016). Both of these assume that BDMIs are, on average, at least partially recessive in the hybrids.

First, under dominance theory, heterogametic hybrid malfunction is explained by BDMIs involved in autosomal-sex chromosome interactions (Turelli and Orr 1995). Whereas hybrids of the homogametic sex will express only those X (or Z)-linked BDMIs that are at least partially dominant, hybrids of the heterogametic sex will express all X (or Z)-linked BDMIs, regardless of dominance. There is empirical evidence for the dominance theory, particularly for inviability loci. Most of these loci have been identified in *Drosophila* (Heikkinen and Lumme 1998; Coyne et al. 2004; Masly and Presgraves 2007), but also many other plant and animal taxa (Salazar et al. 2005; Carling and Brumfield 2008; Brothers and Delph 2010; Demuth et al. 2013).

The faster-X explanation for Haldane’s rule stems from the observation that the X (or Z) chromosome often has a disproportionate impact on hybrid fitness compared to autosomes, a pattern known as the large X-effect (Charlesworth et al. 1987). One explanation for the large X-effect is that new beneficial mutations that are partially recessive will have a faster substitution rate on the X chromosome compared to the autosomes (Charlesworth et al. 1987). This is because on the X chromosome, new recessive alleles are immediately visible to selection in heterogametic individuals. This faster accumulation of substitutions on the X provides more opportunities for BDMIs to arise. Faster-X evolution can lead to Haldane’s rule either via exacerbating the effect of dominance described above or via the fixation of alleles that act in the heterogametic sex only (Coyne and Orr 2004).

A shared feature of dominance and faster-X theories is that the expression of recessive alleles on sex chromosomes in the heterogametic sex results in stronger postzygotic isolation compared to the homogametic sex. All else equal, both mechanisms predict that the rate of evolution of IPI should correlate positively with the extent of hemizygosity. In support of this prediction, *Drosophila* species that have a larger proportion of their genome on the X chromosome evolve IPI more rapidly than species with smaller X chromosomes (Turelli and Begunt 1997). Additionally, taxa with heteromorphic sex chromosomes evolve IPI at lower levels of genetic divergence than taxa with homomorphic or no sex chromosomes (Lima 2014).

Although Haldane’s rule has predominantly been studied in diploid taxa with sex chromosomes, it is also applicable to haplodiploids (Haldane 1922; Koevoets and Beukeboom 2009). In haplodiploids, males develop from unfertilized eggs and are haploid, and females develop from fertilized eggs and are diploid (Normark 2003). Thus, in haplodiploid systems the entire genome is analogous to a sex chromosome. Because hemizygosity is maximized in haplodiploids, dominance and faster-X theory predict that evolution of IPI should be maximized in haplodiploid taxa (Koevoets and Beukeboom 2009).

Although there is some empirical evidence of Haldane’s rule in haplodiploids (Koevoets et al. 2012), there are currently no direct comparisons between the rate of IPI evolution in diploids and haplodiploids. Here, we take advantage of a recent study that surveyed the literature and used linear regression to estimate the relationship between genetic divergence and the strength of IPI for diploid taxa with heteromorphic sex chromosomes (Lima 2014). Using this regression line, we asked whether the observed level of IPI in a haplodiploid species pair exceeds the expected IPI for diploid taxa at a comparable level of genetic divergence, as predicted under both dominance and faster-X theories.

To estimate IPI in a haplodiploid species pair, we focused on a pair of sister species in the pine-sawfly genus *Neodiprion*: *N. pinetum* and *N. lecontei* (Order: Hymenoptera; Family: Diprionidae) (Linnen and Farrell 2008). These species have substantial *extrinsic* postzygotic isolation stemming from oviposition traits (Bendall et al. 2017). Specifically, whereas *N. pinetum* females embed their eggs within the needles of a thin-needled pine species (*Pinus strobus*), *N. lecontei* females deposit their eggs in thicker, more resinous needles in other pine species. While females of each species have oviposition traits well-suited to their respective hosts, hybrid females have maladaptive combinations of oviposition traits that lead to hatching failure: they prefer the thin-needled host, but have traits better suited to thicker, more resinous needles. More generally, this species pair has many morphological and behavioral differences and many fixed genetic differences (genome-wide F_ST_ = 0.6, unpublished data). Overall, given the substantial genetic and phenotypic divergence between this species pair and the complete hemizygosity of haploid males, we expected hybrid males to be sterile or inviable. Shockingly, we found no evidence of IPI. In the discussion, we consider possible explanations for this surprising result, including a frequently overlooked quirk of haplodiploid genetics that may drastically slow the emergence of complete IPI in hybrid haploid males.

## Methods

### Study System Details and Overall Approach

The *N. pinetum* and *N. lecontei* lab lines that were used in this study were derived from larvae collected in the field (Table S1) and propagated in the lab for 1-4 generations following standard lab protocols (Harper et al. 2016; Bendall et al. 2017). We evaluated hybrid female and hybrid male viability and fertility relative to their purebred counterparts according to the crossing scheme illustrated in Figure S1. As is the case in most hymenopterans, unfertilized *Neodiprion* eggs give rise to haploid males, while fertilized eggs give rise to diploid females. Thus, interspecific crosses create hybrid females (“F_1_”) and pure-species males. To obtain hybrid males (“F_2_”), we allowed hybrid females to reproduce. For these experiments we used a combination of mated females (produce both males and females if fertilization is successful) and virgin females (produce all-male colonies). Because females of both species lay their entire egg complement on a single branch terminus and colonies are gregarious throughout development (Coppel and Benjamin 1965), all viability measures were colony-level measurements. Additionally, for our viability estimates, we only used colonies that produced live adults. This enabled us to rule out non-IPI related sources of colony failure, such as lab pathogens, diapause, or extrinsic postzygotic isolation (Coppel and Benjamin 1965; Bendall et al. 2017). Fertility measures were based on the reproductive success of individual adults.

With these data, we evaluated presence/absence of hybrid inviability in each sex and in both directions of the cross. We also evaluated presence/absence of hybrid sterility in both directions of the cross for females and in one direction of the cross for males (due to sample limitations). If we observed any evidence of viability or fertility for a particular cross/sex combination, we considered that combination to lack IPI. These qualitative measures of IPI were comparable to published IPI measures from diploid taxa (Coyne and Orr 1989; Lima 2014).

### IPI in females

To evaluate hybrid female viability, we crossed *N. lecontei* females with *N. pinetum* males (LxP) and vice versa (PxL). We then released each mated female into a mesh cage with two *P. strobus* and two *P. banksiana* seedlings (preferred hosts for *N. pinetum* and *N. lecontei*, respectively). When the female oviposited, we reared the resulting colonies on the host chosen for oviposition. To evaluate hybrid female viability, we calculated the proportion of colonies that had adult females emerge from crosses that had any adult emergence. We did this for both directions of the hybrid cross (PxL *N* = 15, LxP *N* = 23), as well as purebred *N. pinetum* (*N* = 15) and *N. lecontei* (*N* = 20). To determine if female viability differed among cross types, we performed a logistic regression followed by a Tukey’s HSD post-hoc test.

To evaluate hybrid female sterility, we recorded oviposition success (i.e., whether or not a female laid eggs) and, if the female oviposited, the number of eggs laid for four cross types: F_1(PxL)_ hybrid female mated to a *N. pinetum* male (*N* = 41), F_1(LxP)_ hybrid female mated to a *N. lecontei* male (*N* = 32), *N. lecontei* female mated to a *N. lecontei* male (*N* = 124), and *N. pinetum* female mated to a *N. pinetum* male (*N* = 108). All females were placed into choice cages as described above. To remove possible effects of mating, we also evaluated oviposition success and egg number for three types of virgin females (we did not have F_1(PxL)_ available for this experiment): F_1(LxP)_ females (*N* = 35), *N. lecontei* females (*N* = 58), and *N. pinetum* females (*N* = 86). We performed a logistic regression to test if oviposition willingness differed, and an ANOVA to test if egg number differed between female type. For both analyses we used Tukey’s post-hoc tests. We performed separate analyses for virgin and mated females. All statistical analyses were performed in R (3.6.0)

### IPI in males

To evaluate hybrid male viability, we placed mated females of different types into oviposition cages (a combination of “choice” and “no-choice” cages were used) and reared the resulting offspring to adulthood. We estimated male viability for each cross type as the proportion of colonies that had adult male emergence. To generate F_2(LxP)_ hybrid males, we crossed F_1(LxP)_ hybrid females with either *N. lecontei* or *N. pinetum* males (*N* = 32). To generate F_2(PxL)_ hybrid males, we backcrossed F_1(PxL)_ hybrid females to *N. pinetum* males (*N* = 9). These crosses result in backcross females and F_2_ males. For comparison, we also examined male emergence in pure *N. pinetum* (*N* = 34) and *N. lecontei* (*N* = 18) crosses. We performed a logistic regression and a Tukey’s HSD post-hoc test to determine if hybrids had lower rates of male emergence compared to the pure species.

We evaluated hybrid male sterility in one direction of the cross (LxP; due to availability of males). First, we examined sperm motility in *N. pinetum* (*N* = 20), *N. lecontei* (*N* = 47), and F_2(LxP)_ males (*N* = 39). Upon eclosion from cocoons, adult males were stored at 4°C until use to prolong life. In some cases, males were used in mating assays prior to testes dissection, then returned to 4°C for a minimum of 24 hours until further use. Males were warmed to room temperature for a minimum of one hour prior to dissection. From each male, we removed both testes and placed each testis on a siliconized slide in 50 μl of testes buffer (183 mM KCl, 47 mM NaCl, 10 mM Tris-HCl, pH 6.8). After piercing a testis, we imaged the sperm at 40x with a Nikon E800 DIC. *Neodiprion* males have sperm that form bundles. We recorded sperm motility for each male (both testes combined) as no motility (no moving bundles), low motility (0-35% moving bundles), or normal motility (>35% moving bundles), Because mating status did not impact motility, we combined data from unmated and mated males (Chisq= 2.66, p= 0.103). To determine whether hybrid males had reduced sperm motility, we performed a Kruskal-Wallace test, followed by Tukey’s post-hoc tests.

To test whether hybrid males could mate successfully, we used no-choice mating assays. We placed a single *N. lecontei* female in a clear 3.25-oz container with either a *N. lecontei* (*N* = 36) or F_2(LxP)_ hybrid male (*N* = 37) (*N. pinetum* males and females were not available). We observed each pair for 2 hours and recorded whether they mated during that time. To test if mating success differed between *N. lecontei* and hybrid males, we performed a logistic regression. Mating does not indicate that hybrid males produce viable sperm. To evaluate hybrid male fertility, we placed each mated female in a cage with a *P. banksiana* seedling and reared resulting colonies as described above. For all colonies with a F_2_ father that produced adults, we evaluated whether there was successful fertilization by recording the proportion that produced adult females (diploid females indicate successful fertilization).

### Comparing observed IPI in haplodiploids to predicted IPI in heteromorphic diploid taxa

Lima (2014) conducted a meta-analysis of published IPI estimates for taxa with heteromorphic, homomorphic, and no sex chromosomes. Using logistic regression, he calculated the expected level of IPI (with a 95% confidence interval) for a given genetic distance (Nei’s D) for these three categories. If haplodiploidy is analogous to extreme sex chromosome heteromorphy, we predict that sawflies will have higher levels of IPI than diploids with heteromorphic sex chromosomes at the same genetic distance. To test this prediction, we calculated Nei’s D for *N. pinetum* and *N. lecontei* and compared the observed level of IPI for this species pair to expectations derived from the relationship between Nei’s D and IPI in diploid taxa with heteromorphic sex chromosomes (Lima 2014).

To calculate Nei’s D, we used adegenet (Nei 1978; Jombart 2008) with SNP data derived from ddRAD sequencing of 44 *N. lecontei* and 23 *N. pinetum* individuals (data from Bendall *et al.* in prep). The individuals in the genetic dataset were from the same populations that established the lab lines we used to measure IPI. To calculate overall IPI between this species pair, we used the scale from Coyne and Orr (1989), which ranges from 0 (no IPI) to 1 (complete IPI). In brief, each sex that is either completely inviable or infertile in each direction of the cross adds 0.25 to the IPI score. We also calculated sex-specific IPI as in Lima (2014). IPI for each sex could take on three possible values: 0, if the sex was viable and fertile in both directions of the cross; 0.5, if the sex was inviable or infertile in one direction only; and 1, if the sex was inviable or infertile in both directions. With the estimates of IPI and genetic distance, we asked whether our observed IPI fell outside of Lima’s (2014) 95% confidence interval for IPI in heteromorphic taxa for our observed genetic distance. We compared observed to expected IPI for both overall and sex-specific measures of IPI.

## Results

### IPI in females

Interspecific crosses produce just as many colonies with viable female adults as intraspecific crosses (Figure 1A; Chisq = 2.20, *P* = 0.53). These data indicate that hybrid females are viable in both directions of the cross. Hybrid females are also fertile in both directions of the cross. Whether mated or virgin, hybrid females are no less willing to oviposit than non-hybrid females (Figure 1B, Figure S2A, Table S2, Table S3). When mated hybrid females oviposited, they also laid just as many eggs as *N. lecontei* and more eggs than *N. pinetum* (Figure 1C, Figure S2B, Table S2, Table S3). Egg number was similar for all virgin females (*P*=0.053, Table S2). Overall, hybrid females are viable and fertile in both directions of the cross.

**Figure 1.**
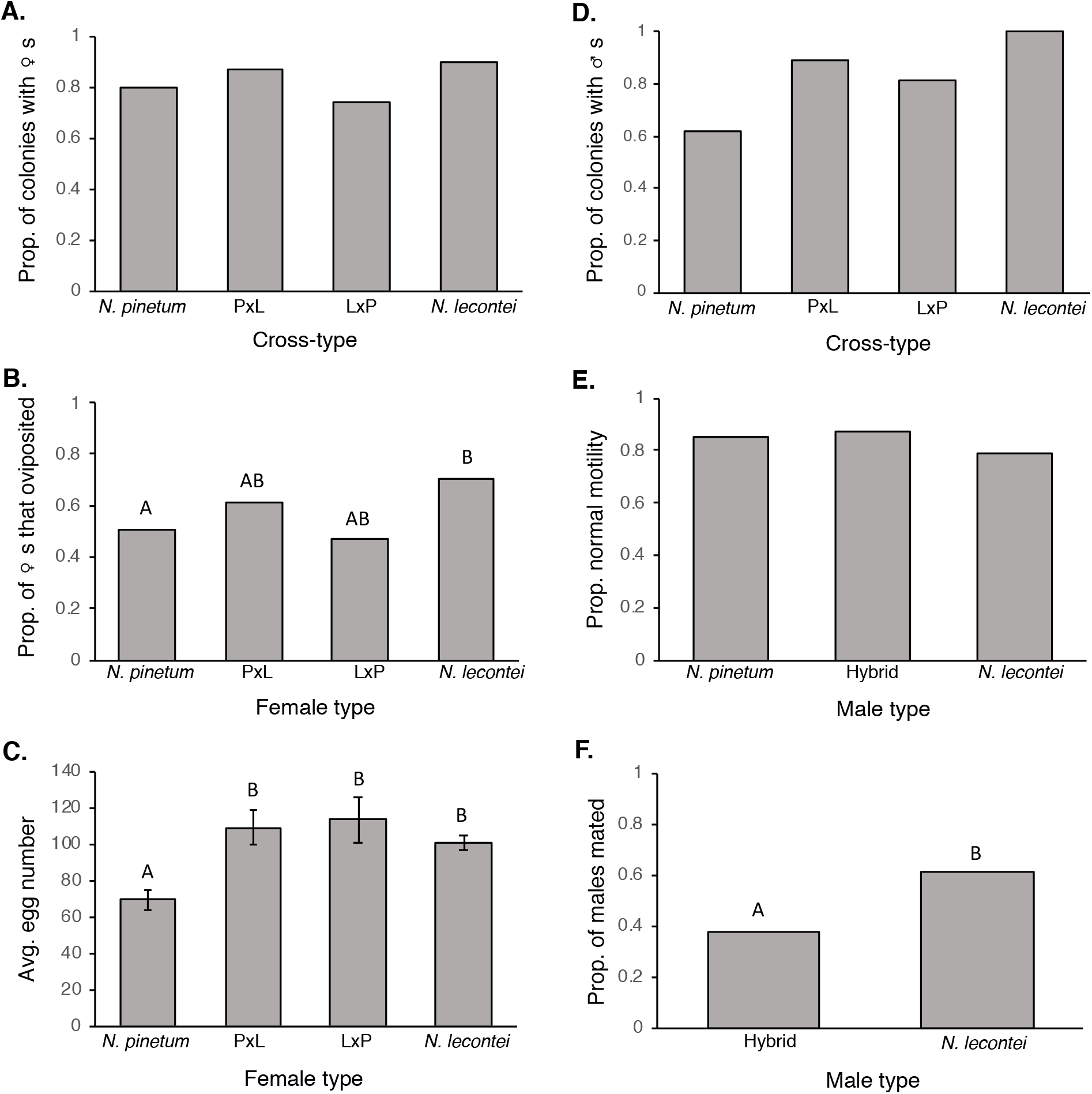
Viability and fertility for hybrid females and hybrid males. **A.** The proportion of colonies with adult females out of all colonies with adult emergence. **B.** The proportion of females that oviposited. **C.** The average number of eggs laid for pure and hybrid females. **D.** The proportion of colonies with adult males out of all colonies with adult emergence. **E.** Proportion of males that had normal sperm motility**. F.** Compared to *N. lecontei* males, hybrid males mate less frequently with *N. lecontei* females. Error bars represent standard error. Different letters denote pairwise comparisons that are significantly different in post-hoc tests; lack of letters indicate that there were no significant differences.

### IPI in males

Hybrid males are viable in both directions of the cross. Although the proportion of colonies that produced adult males varied among cross type (Figure 1D, Chisq = 14.5, *P* = 0.002), hybrid males didn’t have reduced male emergence compared to the pure species (Table S4).

Compared to *N. lecontei*, hybrid F_2(LxP)_ males did not have reduced sperm motility (Figure 1E; Chisq= 1.03, *P*= 0.60). However, *N. lecontei* females were less willing to mate with hybrid males than they were with *N. lecontei* males (Figure 1F; Chisq =3.93, *P* = 0.045). This constitutes a form of *extrinsic* postzygotic isolation (behavioral isolation) in at least one direction of the cross. Nevertheless, hybrid males did mate successfully with some *N. lecontei* females. Of the 10 hybrid-male-fathered colonies that produced adults, 70% produced adult females, indicating that hybrid males are fertile. Overall, hybrid males are viable in both directions of the cross and fertile in one direction (*N. lecontei* female x *N. pinetum* male). The fertility of the reciprocal cross is unknown.

### Genetic distance

Using 21,590 SNPs genotyped in sympatric *N. lecontei* and *N. pinetum* populations, our Nei’s D estimate was 0.36. If we assume that the untested hybrid male type (which differs from the tested hybrid male type only in the mitochondrial genome) was fertile, *N. pinetum* and *N. lecontei* have an IPI score of 0. For a Nei’s D of 0.36, this IPI score is outside of the 95% confidence interval for expected IPI in heteromorphic taxa (Figure 2). However, IPI deviated in the opposite direction of what we predicted: *N. pinetum* and *N. lecontei* have lower IPI than expected given their genetic distance. The individual sexes also had IPI scores of 0. While the male-specific IPI was lower than the 95% confidence interval, the female-specific IPI fell within the 95% confidence interval (which included 0).

**Figure 2.**
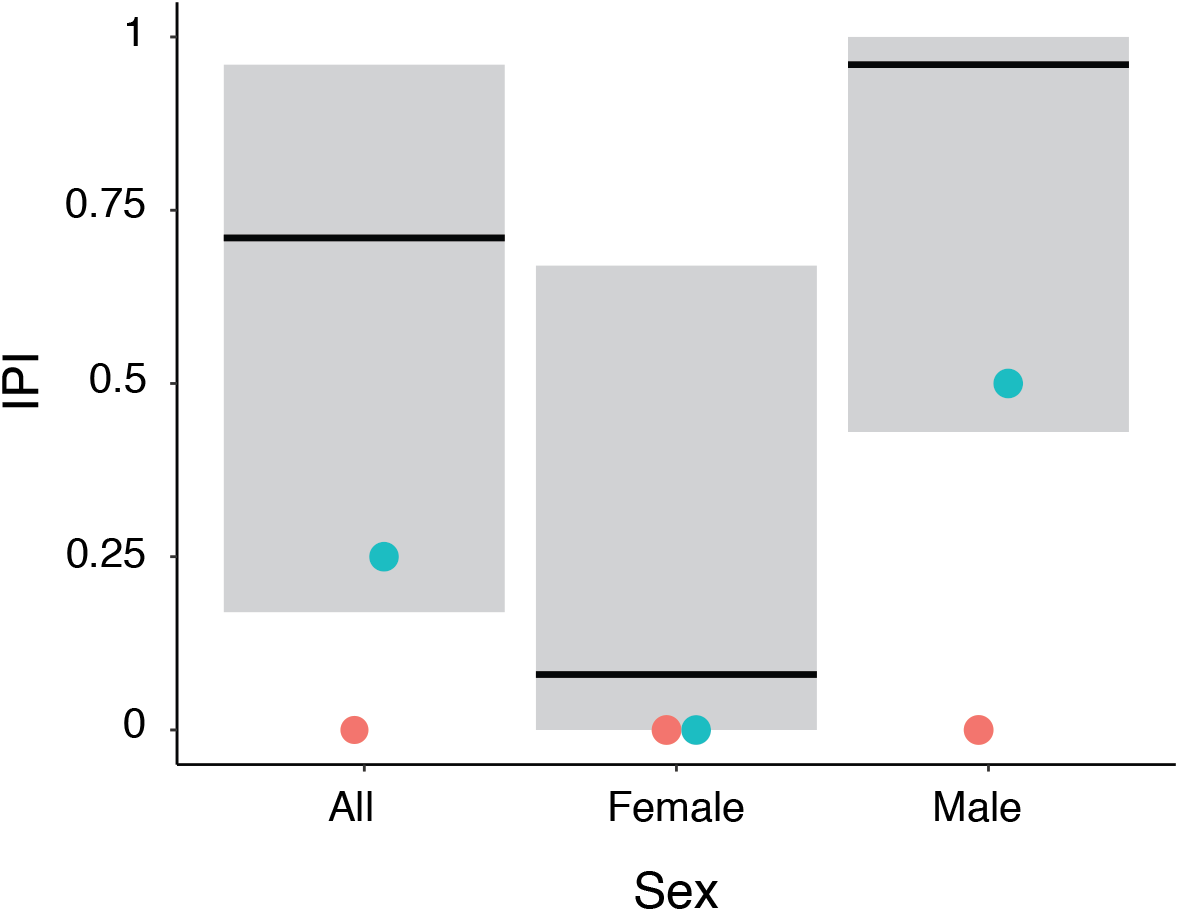
*N. pinetum* and *N. lecontei* have lower isolation than taxa with heteromorphic sex chromosomes at the same genetic distance. The thick line is the expected amount of IPI for taxa with heteromorphic sex chromosomes at Nei’s D of 0.36 (from Lima 2014). The gray shaded area represents the 95% confidence interval. The circles represent the observed isolation between *N. pinetum* and *N. lecontei.* The blue circle is the maximum potential isolation and the pink circle is the observed isolation. Isolation for both sexes combined, females only, and males only are shown.

If we assume instead that the untested hybrid male type was infertile, this would give an overall IPI of 0.25, a male-specific IPI of 0.5, and a female-specific IPI of 0. These IPI estimates, which are the maximum possible IPI for this species pair, did fall within the 95% confidence interval of the heteromorphic taxa. However, these adjusted IPI scores still fell below the heteromorphic regression lines.

## Discussion

Haplodiploids should have higher levels of IPI than diploids for a given genetic distance because all recessive mutations are expressed in the haploid males. We found that *N. lecontei* and *N. pinetum* hybrids are fertile and viable for both sexes, making the IPI score lower than expected given the genetic distance between *N. pinetum* and *N. lecontei*. Here, we consider several possible reasons for the lack of intrinsic isolation in hybrid haploid males.

First, not all genetic mechanisms that have been hypothesized to explain Haldane’s rule rely on the hemizygous nature of sex chromosomes. Although there is significant support for dominance theory and faster-X, mechanisms that rely on chromosomal segregation or the sex-determining properties of the X chromosome may also contribute to Haldane’s rule. These mechanisms will not cause IPI to evolve more rapidly--and may even slow the emergence of IPI--in haplodiploids. For example, under meiotic drive, the sex ratio becomes distorted from 50:50 when drive elements evolve on the X chromosome (McDermott and Noor 2010). Strong negative selection against distorted sex ratios favors suppressors on other chromosomes (autosomal or Y) that restore a balanced sex ratio. This cycle of antagonistic coevolution of drivers and suppressors results in rapid evolution of the X chromosome and increased divergence between species. With increased divergence comes an increased number of incompatibilities. In several *Drosophila* groups meiotic drive has been implicated in hybrid sterility (Hauschteck-Jungen 1990; Tao et al. 2001; Orr and Irving 2005).

In theory, meiotic drive could also produce Haldane’s rule in many haplodiploids. The genetic mechanism underlying haplodiploidy in *Neodiprion* sawflies and many other Hymenoptera is complimentary sex determination, in which sex is determined by heterozygosity at one or more sex-determining loci (Cook 1993; Harper et al. 2016). If an individual is hemizygous (haploid) or homozygous (diploid) at all sex-determining loci, then they are male. Individuals that are heterozygous (diploid) at one or more sex-determining loci are female. Meiotic drive elements can be linked to these sex-determining loci. As drive causes the frequency of an allele at the sex determining locus to increase, the number of homozygous individuals increases. Linked meiotic drive elements would create an unbalanced sex ratio that increases the proportion of diploid males in the population. Diploid males tend to be inviable or sterile, and diploid male production should be strongly selected against (Van Wilgenburg et al. 2006). However, sex-determining loci and linked sites make up a small proportion of the genome. Thus, if meiotic drive is an important source of IPI, haplodiploids should not have faster evolution of IPI, and may evolve IPI more slowly depending on the proportion of the genome that is sex linked.

Although dominance theory has wide support for causing male inviability, there is a lack of support when it comes to sterility (Presgraves 2010). An alternative mechanism to explain the evolution of sterility in the heterogametic sex is incorrect pairing of sex chromosomes during meiosis. Unlike homomorphic chromosomes, which pair by overall homology, heteromorphic sex chromosomes usually match by small stretches of shared sequence, which are often rapidly evolving repeat sequences. These repeat sequences can differ between species, causing hybrid X and Y (or Z and W) chromosomes to be unable to pair or separate properly during meiosis, making the hybrid’s gametes sterile. Since only heteromorphic sex chromosomes require this form of meiotic pairing, this mechanism could explain Haldane’s rule. Chromosome separation failures during meiosis, leading to sterility in hybrids, have recently been reported for mice (Schwahn et al. 2018) mosquitoes (Lang and Sharakhov 2019) yeast (Rogers et al. 2018), and *Drosophila* (Kanippayoor et al. 2020). Because haplodiploids lack sex chromosomes and males do not undergo meiosis to produce gametes, improper pairing of sex chromosomes cannot cause hybrid male sterility in haplodiploids, and hybrid males are therefore fertile.

Even if there is a common underlying genetic mechanism that gives rise to Haldane’s rule across taxa in nature that should give rise to IPI in our system, there could be system-specific reasons why IPI was not observed. For example, the lack of IPI may be due to the divergence history of the species pair we examined, which has ongoing gene flow. If there is sufficient gene flow, deleterious genetic combinations are quickly produced and purged, preventing the evolution of hybrid inviability and sterility (Agrawal et al. 2011). In haplodiploids these deleterious alleles should be purged more quickly (Avery 1984). Whether there is sufficient gene flow in this system to cause selection against IPI remains to be tested.

Alternatively, the low levels of male IPI may be a consequence of haplodiploidy itself. Haplodiploidy may influence the evolution of IPI in more complex ways that depend on the subtleties of the genetic basis of the BDMIs. For example, BDMIs may be formed when there is a mildly deleterious mutation in one species followed by a compensatory mutation in the same species (Kondrashov et al. 2002), as observed in some mitonuclear interactions (Barreto and Burton 2012). When the deleterious mutation is placed into the other genetic background without the compensatory mutation, the hybrid suffers low fitness. Since most mildly deleterious segregating variants in a population are recessive (Simmons and Crow 1977), and all recessive variants are expressed in haplodiploid males, selection will be more efficient at removing these variants from the population, resulting in fewer mildly deleterious mutations in haplodiploids (Avery 1984). If BDMIs formed through compensatory mutations are common, haplodiploids would be expected to evolve IPI more slowly than diploids. This reduced IPI in haplodiploids is only applicable if the dominance theory underlies sterility, since faster-X theory specifically deals with positively selected recessive alleles. To rigorously test this idea, the specific mutations involved in BDMI and their fitness effects in the original population must be known.

Finally, the haplodiploid inheritance mechanism may account for the absence of complete inviability and sterility in hybrid males. To illustrate why, we propose a verbal model describing the effects of haplodiploid inheritance. We consider a simple two-locus BDMI in which there is one locus on the autosome and a second locus on the X chromosome that interact to cause the incompatibility (Figure 3A). In one species, a derived co-dominant autosomal mutation fixes, and in the other species a recessive mutation fixes on the X chromosome. When these species hybridize, hybrids are sterile or inviable when one copy of the derived autosomal allele and only the derived X allele is present in the hybrid, as would be observed in the heterogametic sex. Only one direction of the cross will experience IPI under this simple model.

**Figure 3.**
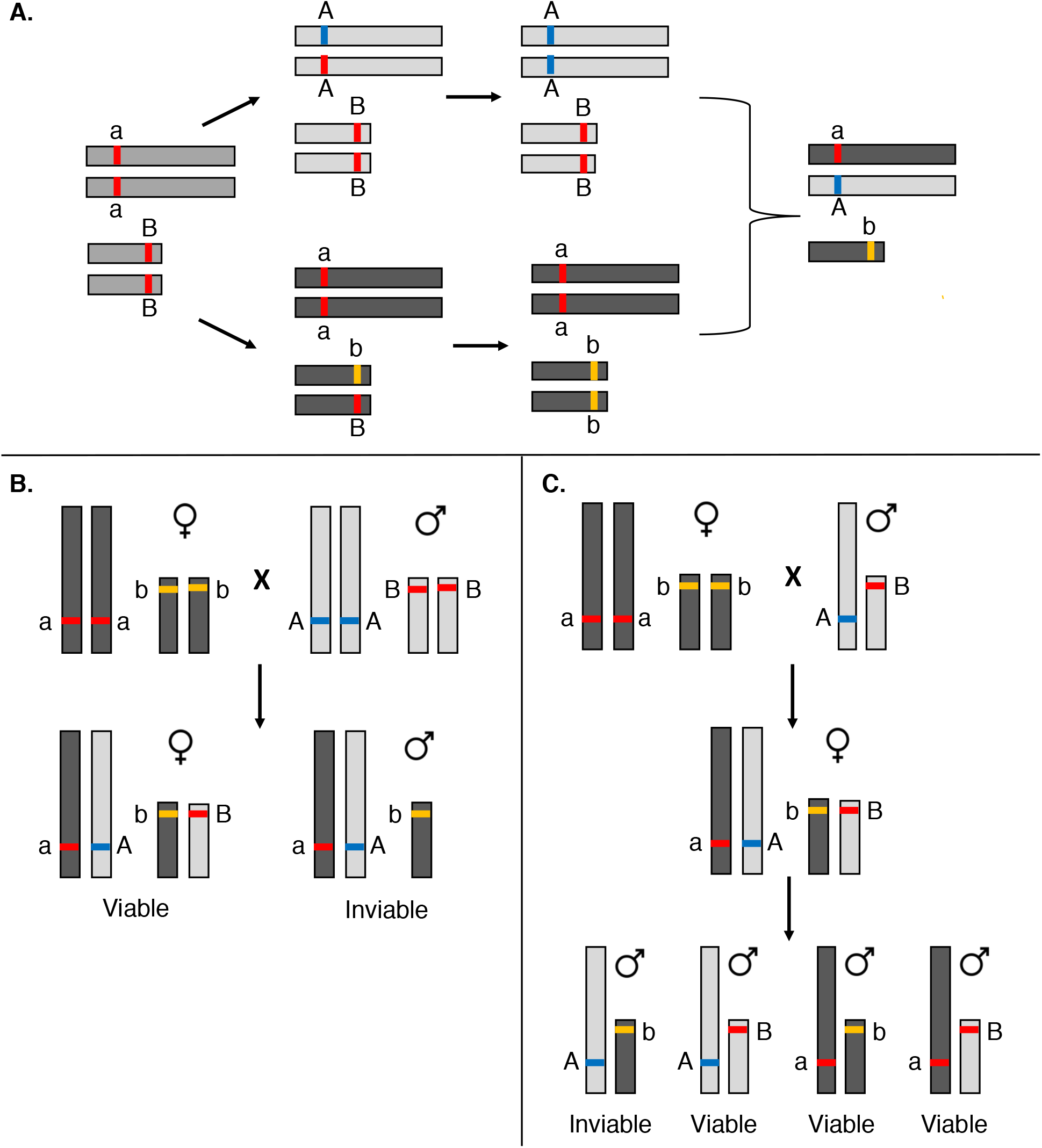
A two-locus model of IPI in haplodiploids and diploids consistent with the dominance model. **A.** The evolution of a two-locus BDMI in diploids with incompatible loci on an autosome (long rectangle) and X chromosome (short rectangle). The red uppercase alleles are ancestral. The X-linked derived allele (yellow b) is recessive and the autosomal derived allele (blue a) is at least codominant. In hybrids, an individual with at least one A allele and homozygous or hemizygous for the b allele, such as the male shown at the right, will be inviable or sterile. Only one direction of the cross will experience IPI under this model **B.** In diploids, hybridization leads to viable females and inviable males. All males have the same genotype that include the incompatibility. **C.** In haplodiploids, hybrid females are viable. Hybrid males aren’t formed until the second generation. Only 25% of the males will have the incompatibility and be inviable.

For the direction of the cross with IPI, all diploid hybrid males have one set of autosomal chromosomes from each species and the X from the maternal species causing all males to have the incompatibility (Figure 3B). The females will be viable and fertile, since the incompatible X locus is heterozygous. In haplodiploids, the incompatibilities would be on two different autosomes since they do not have sex chromosomes (Figure 3C). A F_1_ female has the same genotype as the diploid female and is viable and fertile. However, there is no true F_1_ hybrid male. Instead, the first generation of hybrid males (F_2_) are the offspring of hybrid females, allowing for recombination to occur before hybrid males are formed. Unlike diploids, not all males will have all incompatibilities. Instead, only 25% of the males will have the derived allele at both loci and will be inviable or infertile in this two-locus model. Although this model is more simplistic than many BDMIs in nature, it shows how recombination in F_1_ females allows viable allelic combinations to be formed in hybrid males. The exact effect of haplodiploidy inheritance on the evolution of IPI will depend on the genetic architecture of the BDMIs (e.g. effect size, dominance, genomic locations), and further modeling is necessary.

We have proposed several explanations for why we detected lower IPI than expected under dominance theory and faster-X. These explanations fall into three categories 1) system specific effects 2) inheritance mechanism of haplodiploids, and 3) alternative mechanisms for Haldane’s rule. Meiotic drive, pairing of sex chromosomes during meiosis, and dominance theory formed through compensatory mutations are non-mutually exclusive mechanisms that may all be important drivers of the evolution of BDMIs and Haldane’s rule. Importantly, haplodiploids can be used to distinguish between these different mechanisms because they have different predictions for the level of IPI in haplodiploids compared to diploids (Table 1). To tease apart these different possibilities, quantitative measures of sterility and inviability from many haplodiploid and diploid taxa are needed. The influence of system-specific effects such as interspecific gene flow and species population sizes should also be examined. Biologists have been trying to understand the mechanism underlying Haldane’s rule for almost a century. Haplodiploids have been an underutilized resource in this search and have the potential to provide novel insight into the underlying basis of this phenomenon.

**Table 1.**
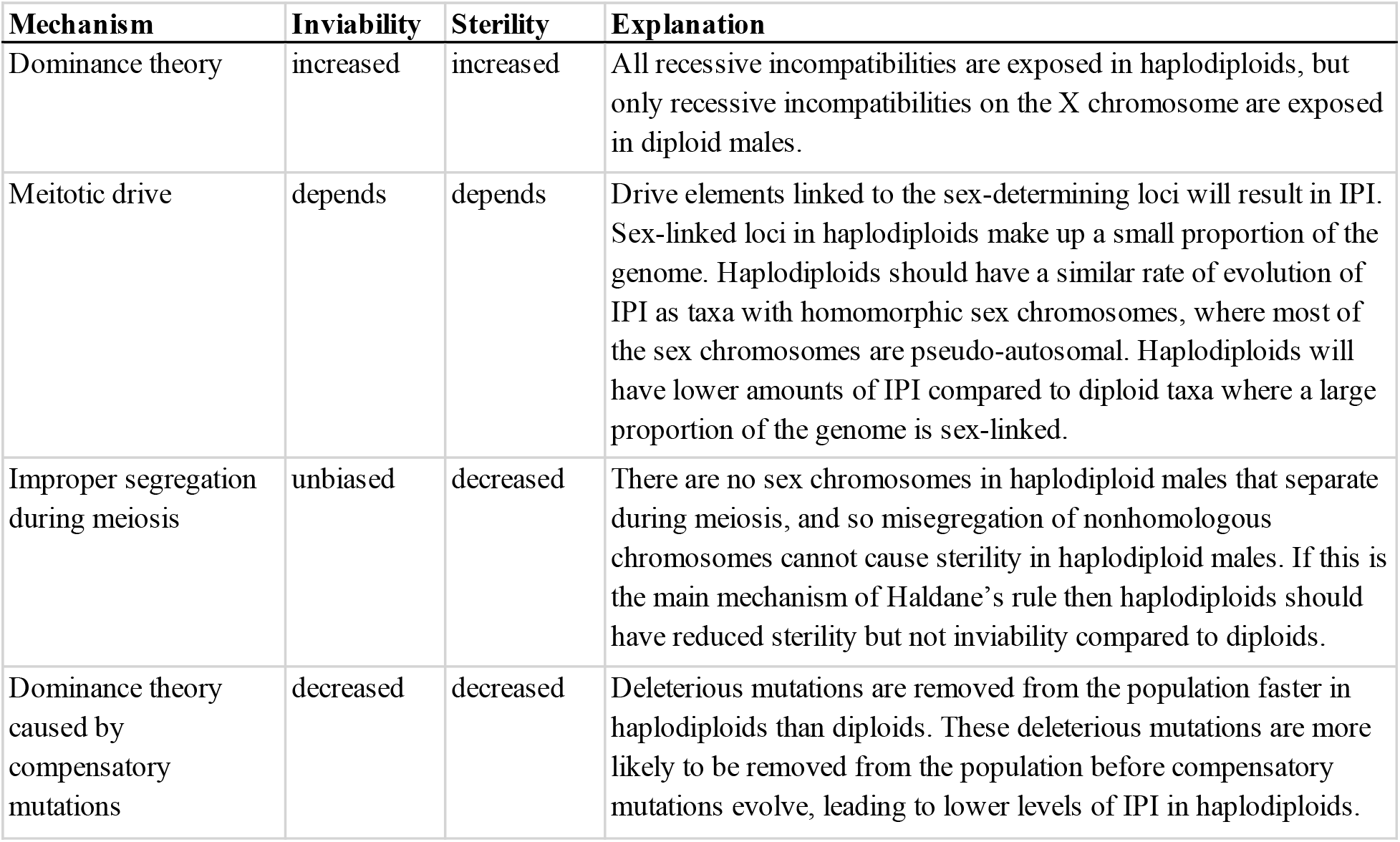
Proposed mechanisms for Haldane’s rule and predictions regarding how haplodiploids should differ from diploids in the rate of evolution of IPI.

| **Mechanism** | **Inviability** | **Sterility** | **Explanation** |
| --- | --- | --- | --- |
| Dominance theory | increased | increased | All recessive incompatibilities are exposed in haplodiploids, but only recessive incompatibilities on the X chromosome are exposed in diploid males. |
| Meiotic drive | depends | depends | Drive elements linked to the sex-determining loci will result in IPI. Sex-linked loci in haplodiploids make up a small proportion of the genome. Haplodiploids should have a similar rate of evolution of IPI as taxa with homomorphic sex chromosomes, where most of the sex chromosomes are pseudo-autosomal. Haplodiploids will have lower amounts of IPI compared to diploid taxa where a large proportion of the genome is sex-linked. |
| Improper segregation during meiosis | unbiased | decreased | There are no sex chromosomes in haplodiploid males that separate during meiosis, and so missegregation of nonhomologous chromosomes cannot cause sterility in haplodiploid males. If this is the main mechanism of Haldane's rule then haplodiploids should have reduced sterility but not inviability compared to diploids. |
| Dominance theory caused by compensatory mutations | decreased | decreased | Deleterious mutations are removed from the population faster in haplodiploids than diploids. These deleterious mutations are more likely to be removed from the population before compensatory mutations evolve, leading to lower levels of IPI in haplodiploids. |

## Acknowledgements

We would like to thank members of the Linnen lab for help with collecting and rearing of the sawflies. For advice on testes dissection, we thank Rachelle Kanippayoor. For help with sperm microscopy, we thank Doug Harrison. For useful discussion, we thank Harmit Malik. Funding was provided by the National Science Foundation (DEB-1257739 & CAREER-175096, both to CRL).

## Literature cited

Agrawal, A., J. Feder, and P. Nosil. 2011. Ecological divergence and the origins of intrinsic postmating isolation with gene flow. Int. J. Ecol. 435357.

Avery, P. J. 1984. The population genetics of haplo-diploids and X-linked genes. Genet. Res., Camb 44:321–341.

Barreto, F. S., and R. S. Burton. 2012. Evidence for compensatory evolution of ribosomal proteins in response to rapid divergence of mitochondrial rRNA. Mol. Biol. Evol 30:310–314.

Bateson, W. 1909. Heredity and variation in modern lights. Pp. 85–101 in A. C. Seward, ed. Danvin and Modern Science. Cambridge University Press, Cambridge.

Bendall, E. E., R. K. Bagley, V. C. Sousa, and C. R. Linnen. n.d. Haplodiploidy leads to greater and more variable genomice divergence.

Bendall, E. E., K. L. Vertacnik, and C. R. Linnen. 2017. Oviposition traits generate extrinsic postzygotic isolation between two pine sawfly species. BMC Evol. Biol. 17.

Brothers, A. N., and L. F. Delph. 2010. Haldane’s rule is extended to plants with sex chromosomes. Evolution (N. Y). 64:3643–3648.

Carling, M. D., and R. T. Brumfield. 2008. Haldane’s rule in an avian system using cline theory and divergence population genetics to test for differential introgression of mitochondrial, autosomal, and sex-linked loci across the Passerina bunting hybrid Zone. Evolution (N. Y). 62:2600–2615.

Charlesworth, B., J. A. Coyne, and N. H. Barton. 1987. The relative rates of evolution of sex chromosomes and autosomes.

Cook, J. M. 1993. Sex determination in the Hymenoptera: a review of models and evidence.

Coppel, H., and D. Benjamin. 1965. Bionomics of the-nearctic pine-feeding diprionids. Annu. Rev. Entomol. 10:69–96.

Coyne, J. A., and H. A. Orr. 1997. “Patterns of speciation in Drosophila” revisited. Evolution (N. Y). 51:295–303.

Coyne, J., S. Elwyn, S. Kim, A. L.-G. Research, and U. 2004. 2004. Genetic studies of two sister species in the Drosophila melanogaster subgroup, D. yakuba and D. santomea. Genet. Res. Camb. 84:11–26.

Coyne, J., and H. Orr. 1989. Patterns of Speciation in Drosophila. Evolution (N. Y). 43:362–381.

Coyne, J., and H. Orr. 2004. Speciation. Sinauer Associates, Sunderland, MA.

Delph, L. F., and J. P. Demuth. 2016. Haldane’s rule: Genetic bases and their empirical support. J. Hered. 107:383–391.

Demuth, J. P., R. J. Flanagan, and L. F. Delph. 2013. Genetic architecture of isolation between two species of Silene with sex chromosomes and Haldane’s rule. Evolution (N. Y). 68:332–342.

Dobzhansky, T. 1937. Genetics and the origin of species. 3rd ed. Columbia University Press, New York.

Dobzhansky, T. 1951. Genetics and the Origin of Species. 3rd ed. Columbia University Press, New York.

Haldane, J. B. S. 1922. Sex ration and unisexual sterility in hybrid animals. J. Genet. 12:101–109.

Harper, K., R. Bagley, K. Thompson, and C. Linnen. 2016. Complementary sex determination, inbreeding depression and inbreeding avoidance in a gregarious sawfly. Heredity (Edinb). 117:326–335.

Hauschteck-Jungen, E. 1990. Postmating reproductive isolation and modification of the ‘sex ratio’ trait in Drosophila subobscura induced by the sex chromosome gene arrangement. Genetica 83:31–44. Kluwer Academic Publishers.

Heikkinen, E., and J. Lumme. 1998. The Y chromosomes of Drosophila lummei and D. novamexicana differ in fertility factors. Heredity (Edinb). 81:505–513. Nature Publishing Group.

Jombart, T. 2008. adegenet: a R package for the multivariate analysis of genetic markers. Bioinformatics 24:1403–1405. Narnia.

Kanippayoor, R. L., J. H. M. Alpern, and A. J. Moehring. 2020. A common suite of cellular abnormalities and spermatogenetic errors in sterile hybrid males in Drosophila. Proc. R. Soc. B In press.

Koevoets, T., and L. W. Beukeboom. 2009. Genetics of postzygotic isolation and Haldane’s rule in haplodiploids. Heredity (Edinb). 102:16–23. Nature Publishing Group.

Koevoets, T., O. Niehuis, L. van de Zande, and L. W. Beukeboom. 2012. Hybrid incompatibilities in the parasitic wasp genus Nasonia: negative effects of hemizygosity and the identification of transmission ratio distortion loci. Heredity (Edinb). 108:302–311. Nature Publishing Group.

Kondrashov, A. S., S. Sunyaev, and F. A. Kondrashov. 2002. Dobzhansky-Muller incompatibilities in protein evolution. Proc. Natl. Acad. Sci. U. S. A. 99:14878–83. National Academy of Sciences.

Lang, J., and I. Sharakhov. 2019. Premeiotic and meiotic failures lead to hybrid male sterility in the Anopheles gambiae complex. Proc. R. Soc. B 286:20191080.

Lima, T. G. 2014. Higher levels of sex chromosome heteromorphism are associated with markedly stronger reproductive isolation. Nat. Commun. 5:4743.

Linnen, C. R., and B. D. Farrell. 2008. Comparison of methods for species-tree inference in the sawfly genus Neodiprion (Hymenoptera: Diprionidae). Syst. Biol. 57:876–90.

Masly, J. P., and D. C. Presgraves. 2007. High-resolution genome-wide dissection of the two rules of speciation in Drosophila. PLoS Biol. 5:e243.

McDermott, S. R., and M. A. F. Noor. 2010. The role of meiotic drive in hybrid male sterility. Phil. Trans. R. Soc. B 365:1265–1272.

Moehring, A., A. Llopart, S. Elwyn, … J. C.-, and U. 2006. 2006. The genetic basis of postzygotic reproductive isolation between Drosophila santomea and D. yakuba due to hybrid male sterility. Genetics 173:225–233.

Muller, H. J. 1942. Isolating mechanisms, evolution and temperature. Biol. Symp. 6:71–125.

Nei, M. 1978. Estimation of average heterozygosity and genetic distance from a small number of individuals. Genetics 89:583–590.

Normark, B. B. 2003. The evolution of alternative genetic systems in insects. Annu. Rev. Entomol 48:397–423.

Orr, H. A., and S. Irving. 2005. Segregation distortion in hybrids between the Bogota and USA subspecies of Drosophila pseudoobscura. Genetics 169:671–682.

Presgraves, D. C. 2010. The molecular evolutionary basis of species formation. Nat. Rev. Genet. 11:175–80. Nature Publishing Group.

Rogers, D., E. McConnell, J. Ono, and D. Greig. 2018. Spore-autonomous fluorescent protein expression identifies meiotic chromosome mis-segregation as the principal cause of hybrid sterility in yeast. PLoS Biol. 16:e2005066.

Salazar, C. A., C. D. Jiggins, C. F. Arias, A. Tobler, E. Bermingham, and M. Linares. 2005. Hybrid incompatibility is consistent with a hybrid origin of Heliconius heurippa Hewitson from its close relatives, Heliconius cydno Doubleday and Heliconius melpomene Linnaeus. J. Evol. Biol. 18:247–256.

Schilthuizen, M., M. C. W. G. Giesbers, and L. W. Beukeboom. 2011. Haldane’s rule in the 21st century. Heredity (Edinb). 107:95–102. Nature Publishing Group.

Schwahn, D., R. Wang, M. White, and B. Payseur. 2018. Genetic dissection of hybrid male sterility across stages of spermatogenesis. Genetics 210:1453–1465.

Simmons, M. J., and J. F. Crow. 1977. Mutations affecting fitness in Drosophila populations. Ann. Rev. Genet. 11:49–78.

Tao, Y., D. L. Hartl, and C. C. Laurie. 2001. Sex-ratio segregation distortion associated with reproductive isolation in Drosophila. Proc Natl. Acad. Sci. U.S.A 98:13183–13188.

Turelli, M., and D. J. Begunt. 1997. Haldane’s Rule and X-chromosome Size in Drosophila. Genetics 147:1799–1815.

Turelli, M., and H. A. Orr. 1995. The dominance theory of Haldane’s rule. Genetics 140:389–402.

Van Wilgenburg, E., G. Driessen, and L. W. Beukeboom. 2006. Single locus complementary sex determination in Hymenoptera: an “unintelligent” design? Front. Zool. 3:1.

