## Supplemental Materials for "Lack of intrinsic postzygotic isolation in haplodiploid male hybrids despite high genetic distance"

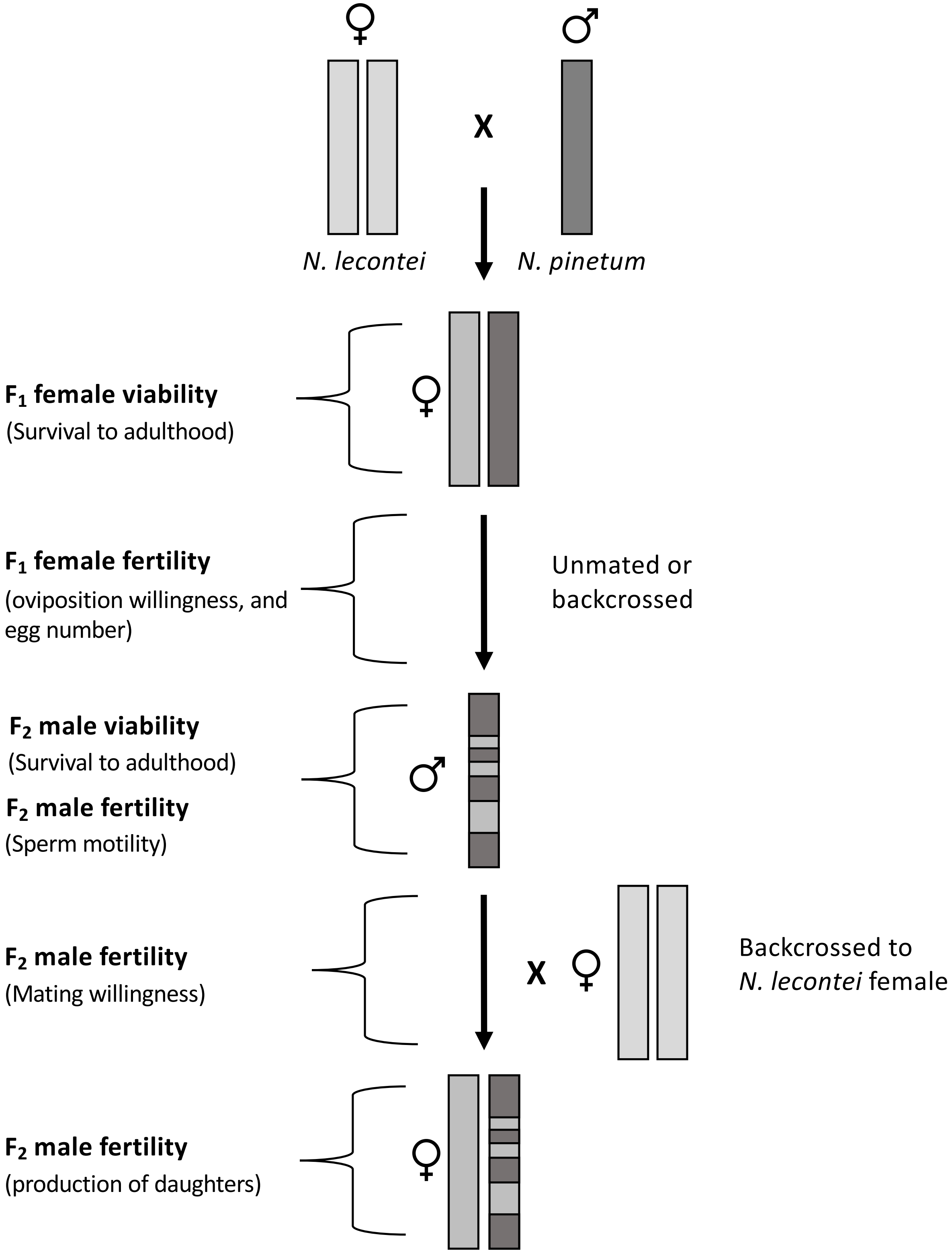

Figure S1. Diagram of one direction of the crosses illustrating when each set of reproductive isolation data was collected. The reciprocal cross was also performed for $F_1$ female viability and fertility and $F_2$ male viability. The rectangles represent the genotype of the individual. Light grey is *N. lecontei* and dark grey is *N. pinetum* genetic material . The haploids (single rectangle) are males and the diploids (2 rectangles) are female.

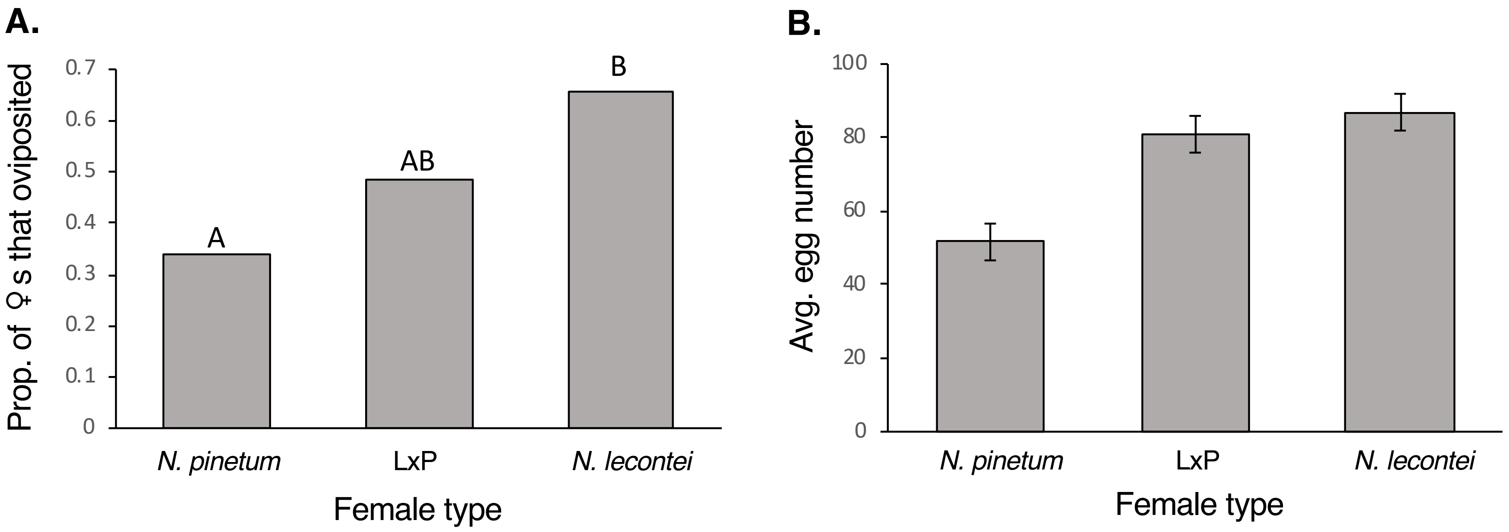

Figure S2. Virgin female fertility. A**.** The proportion of females that oviposited. **B.** The average number of eggs laid for pure and hybrid females. Error bars represent SE. Different letters denote pairwise comparisons that are statistically significantly different in post-hoc tests; lack of letters indicate that there were no significant differences.

Table S1. Sampling locations for *N. pinetum* and *N. lecontei* used in fertility and viability estimates.

| | Population | Female viability | Female fertility | Male viability | Male sperm motility | Male mating success | Male fertility - female production | State | Lattitude | Longitude |
|---|---|---|---|---|---|---|---|---|---|---|
| *N. lecontei* | | | | | | | | | | |
| | Arboretum | X | X | X | X | X | X | KY | 38.014 | -84.504 |
| | Crossville, TN | | X | X | | X | | TN | 35.980 | -85.015 |
| | Spooner | | X | | | | | WI | 45.822 | -91.888 |
| | Grayling, MI | X | X | X | | | | MI | 44.657 | -84.696 |
| | High St. | | | | | X | X | KY | 38.044 | -84.497 |
| | Tates Creek | | | | X | | | KY | 37.970 | -84.511 |
| | Clay's Mill | | | | X | | | KY | 37.984 | -84.559 |
| *N. pinetum* | | | | | | | | | | |
| | Ecton Park | | X | | | | | KY | 38.016 | -84.490 |
| | Regency Rd | X | X | X | | | | KY | 38.003 | -84.525 |
| | Man 'O War | | X | | | | | KY | 37.971 | -84.498 |
| | Georgetown South | X | X | X | X | X | X | KY | 38.249 | -84.549 |
| | Georgetown North | X | X | | X | X | | KY | 38.248 | -84.545 |
| | Cardinal Run | X | X | X | X | | | KY | 38.032 | -84.565 |
| | Belleau Woods | X | X | X | | | | KY | 37.973 | -84.500 |
| | Crossville, TN | X | X | X | | | | TN | 35.980 | -85.015 |
| | McConnel Springs | X | X | X | X | X | X | KY | 38.056 | -84.528 |
| | Starshoot Pkwy | X | X | X | X | X | X | KY | 38.025 | -84.423 |
| | Walton | | X | | | | | KY | 38.857 | -84.618 |
| | Florence | | X | | | | | KY | 39.008 | -84.650 |
| | Waverly | X | | | | | | KY | 37.989 | -84.574 |

Table S2. Statistics for mated and virgin female fertility. Single step adjusted P-values are reported.

| Fertility Measurement | test-statistic | test-statistic value | df | P |
|---|---|---|---|---|
| Mated oviposition willingness | Chisq | 11.415 | 3 | 9.68E-03 |
| Mated egg number | F | 8.981 | 3 | 1.51E-05 |
| Virgin oviposition willingness | Chisq | 14.316 | 2 | 7.78E-04 |
| Virgin egg number | F | 3.113 | 2 | 0.0537 |

Table S3. Tukey's HSD post-hoc tests for mated and virgin female fertility. Single step adjusted P-values are reported.

| | Mated oviposition willingness | | Mated egg number | | Virgin oviposition willingness | |
|---|---|---|---|---|---|---|
| Comparison | Z value | P | t value | P | Z value | P |
| *N. pinetum* vs. PxL | -1.10 | 0.685 | 4.04 | <0.001 | NA | NA |
| *N. pinetum* vs LxP | -0.402 | 0.977 | 3.78 | 1.27E-03 | 1.52 | 0.281 |
| *N. pinetum* vs. *N. leconte* i | -2.98 | 0.0145 | -4.29 | <0.001 | -3.68 | <0.001 |
| PxL vs. LxP | 1.20 | 0.621 | -0.355 | 0.984 | NA | NA |
| PxL vs *N. lecontei* | -1.09 | 0.690 | 0.765 | 0.96 | NA | NA |
| LxP vs. *N. lecontei* | -2.42 | 0.0699 | 1.19 | 0.622 | -1.60 | 0.244 |

Table S4. Tukey's HSD post-hoc tests for male viability (colonies that had adult emergence). Single step adjusted P-values are reported.

| Comparison | Z | P |
|---|---|---|
| *N. pinetum* vs. PxL | -1.43 | 0.416 |
| *N. pinetum* vs LxP | -0.868 | 0.784 |
| *N. pinetum* vs. *N. lecontei* | -0.012 | 1 |
| PxL vs. LxP | 1.01 | 0.696 |
| PxL vs *N. lecontei* | -0.011 | 1 |
| LxP vs. *N. lecontei* | -0.011 | 1 |
